# Single molecule studies of the native hair cell mechanosensory transduction complex

**DOI:** 10.1101/2023.12.11.571162

**Authors:** Sarah Clark, Jaba Mitra, Johannes Elferich, April Goehring, Jingpeng Ge, Taekjip Ha, Eric Gouaux

## Abstract

Hearing and balance rely on the conversion of a mechanical stimulus into an electrical signal, a process known as mechanosensory transduction (MT). In vertebrates, this process is accomplished by an MT complex that is located in hair cells of the inner ear. While the past three decades of research have identified many subunits that are important for MT and revealed interactions between these subunits, the composition and organization of a functional complex remains unknown. The major challenge associated with studying the MT complex is its extremely low abundance in hair cells; current estimates of MT complex quantity range from 3-60 attomoles per cochlea or utricle, well below the detection limit of most biochemical assays that are used to characterize macromolecular complexes. Here we describe the optimization of two single molecule assays, single molecule pull-down (SiMPull) and single molecule array (SiMoA), to study the composition and quantity of native mouse MT complexes. We demonstrate that these assays are capable of detecting and quantifying low attomoles of the native MT subunits protocadherin-15 (PCDH15) and lipoma HMGIC fusion partner-like protein 5 (LHFPL5). Our results illuminate the stoichiometry of PCDH15- and LHFPL5-containing complexes and establish SiMPull and SiMoA as productive methods for probing the abundance, composition, and arrangement of subunits in the native MT complex.

**Impact Statement:** In the present work, the authors develop and employ single molecule methods to detect, characterize, and quantitate attomole quantities of the hair cell mechanosensory transduction complex.

## Introduction

The vertebrate sensations of hearing and balance are critical for everyday life, yet the molecular basis of these sensations remains largely unresolved. This is due to the challenges associated with studying the mechanosensory transduction (MT) complex, the protein machinery that is responsible for converting the mechanical stimulus associated with a sound or fluid movement into an electrical signal that is processed by the brain ^1^. Numerous genetic and biochemical studies have identified MT subunits that are important for mechanosensory transduction, including the tip-link proteins, protocadherin-15 (PCDH15) and cadherin-23 (CDH23), which connect adjacent stereocilia and transduce force associated with stereocilia movement into ion channel opening ^2,3^. Transmembrane channel-like proteins 1 and 2 (TMC1/2) are the likely pore-forming subunits of the ion channel and are located at the lower end of the tip-link ^4–6^. Lipoma HMGIC fusion partner-like protein 5 (LHFPL5, also known as TMHS) ^7–9^, transmembrane inner ear protein (TMIE) ^10–12^, and calcium and integrin-binding proteins 2 and 3 (CIB2/3) ^13,14^ are predicted to assemble with TMC1/2 and PCDH15 to form a functional MT complex. Additional proteins, including transmembrane O-methyltransferase (TOMT) ^15,16^, whirlin ^17,18^, and myosin XVa ^19,20^, also play an important role in hearing and possibly interact with MT subunits at different stages of complex assembly.

Structural and biochemical studies have shed light on interactions between some of these subunits and their probable organization in the vertebrate MT complex. Recombinantly-expressed PCDH15 binds to LHFPL5 and CDH23, and their respective protein complexes have been characterized by cryo-EM ^9^ and x-ray crystallography ^21^. Peptides of TMC1 form a complex with CIB2 and CIB3 ^14,22^, and mutagenesis experiments in mouse hair cells suggest that TMIE directly interacts with TMC1 ^12^. In accordance with these studies, the recently elucidated structures of the native *C. elegans* TMC-1 and TMC-2 complexes revealed a dimeric complex that is composed of two copies of TMC-1/2, TMIE, and CIB2/3, the latter of which is known as CALM-1 in *C. elegans* ^23^. Further, while numerous immunoprecipitation and yeast two-hybrid experiments have suggested interactions between MT subunits ^10,12,15,24,25^, these experiments have been hampered by the inability to express biochemically well-behaved TMC1 and TMC2 proteins. TMC1 and TMC2 do not migrate to the plasma membrane in heterologous cell lines ^26^, preventing reconstitution of the vertebrate MT complex. To elucidate the composition and organization of the vertebrate MT complex, it is therefore necessary to use native tissue.

A well-known hurdle in studying the vertebrate MT complex is its extremely low abundance. On the one hand, electrophysiological experiments of mouse hair cells indicate that there are approximately two functional MT channels per stereocilium ^27–29^, which amounts to about three attomoles of MT complex per mouse cochlea. On the other hand, photobleaching experiments report an average of 7.1 TMC1 molecules per stereocilia in the inner hair cells of P4 aged mice and the number of TMC1 molecules varies from ∼8 to 20 depending on the location in the cochlea ^30^. Electron tomography images of labeled PCDH15 on native stereocilia derived from P6-P9 aged mice indicate there are anywhere from 2, to more than 5, PCDH15 dimers per tip, plus many more PCDH15 molecules that are localized on the stereocilia shaft as lateral links ^31^. Quantification and characterization of attomole quantities of protein has been hindered by the available techniques. Biochemical methods that are commonly used to analyze macromolecular complexes, such as western blots, mass spectrometry, or enzyme-linked immunosorbent assays (ELISAs) require femtomoles of material. Ultrasensitive ELISA ^32^ and immuno-PCR ^33,34^ techniques developed in recent years can detect low attomole quantities of protein, but these assays are limited in their ability to accurately measure protein quantity or assess interactions between proteins and their stoichiometry.

Single molecule studies offer many advantages in the study of low abundance macromolecular complexes ^35^ because they allow for the capture of small amounts of material directly from tissue extracts. Here we describe the optimization of two single molecule assays, single molecule pull-down (SiMPull) ^35^ and single molecule array (SiMoA) ^36^, to study the composition and quantity of native mouse MT complexes, demonstrating that we can detect single digit attomoles of PCDH15- and LHFPL5-containing complexes. Our results not only shed light on the composition of the native mouse MT complex, but also establish techniques to quantify low abundance complexes and to establish the stoichiometry of MT subunits.

## Results

### Development of an ultrasensitive SiMPull assay using recombinant PCDH15/LHFPL5

Due to the extremely low abundance of the MT complex in mouse cochlea, we sought to develop an ultrasensitive assay that is capable of detecting attomoles of protein. We employed single molecule pulldown (SiMPull), a technique that combines a conventional pulldown assay with total internal reflection microscopy (TIRF) to examine individual complexes from cell or tissue extracts ^35,37^. The first step of this assay is to capture MT complexes from a cell extract using a monoclonal antibody (mAb) immobilized on a coverslip. Next, a mAb or antibody fragment (Fab) that is linked to a fluorophore is used to detect the complex. Subunit interactions are assessed through colocalization experiments, wherein multiple mAbs/Fabs are labeled with different fluorophores and applied to the slide simultaneously. Subunit stoichiometry is measured by labeling a mAb/Fab with a single GFP or YFP molecule and photobleaching the imaging area. High affinity antibodies are therefore a key component of a successful SiMPull assay. We generated antibodies directed against different regions of PCDH15 by immunizing mice and rabbits with constructs of recombinantly expressed PCDH15 and the PCDH15/LHFPL5 complex (Fig 1a). We obtained two antibodies with exceptional binding kinetics that recognize native PCDH15: 8D1, which binds to the EC11 domain of PCDH15, and 39G7, which recognizes the EC3 domain of PCDH15. 8D1 and 39G7 both bind PCDH15 with a sub-nanomolar K_D_ (Fig. 1b, Supp. Fig. 1) and 8D1 exhibits a remarkably fast on rate of 7 x 10^5^ M^-1^s^-1^. Immunostaining of wild-type (WT) mouse cochlea with these antibodies produces robust fluorescence that is localized to stereocilia tips (Fig. 1c). The 39G7 mAb was recently employed in cryo-electron tomography experiments to elucidate the molecular structures and conformational states of PCDH15 on stereocilia ^31^. Further, we characterized an anti-LHFPL5 antibody and found that it binds to LHFPL5 with ∼1 nM affinity and similarly stains the tips of stereocilia in WT mice (Fig. 1b,c). Determination of the amino acid sequence of the variable domains of these three antibodies allowed us to generate recombinantly expressed Fab constructs, protein constructs that facilitate single molecule photobleaching experiments.

**Figure 1:**
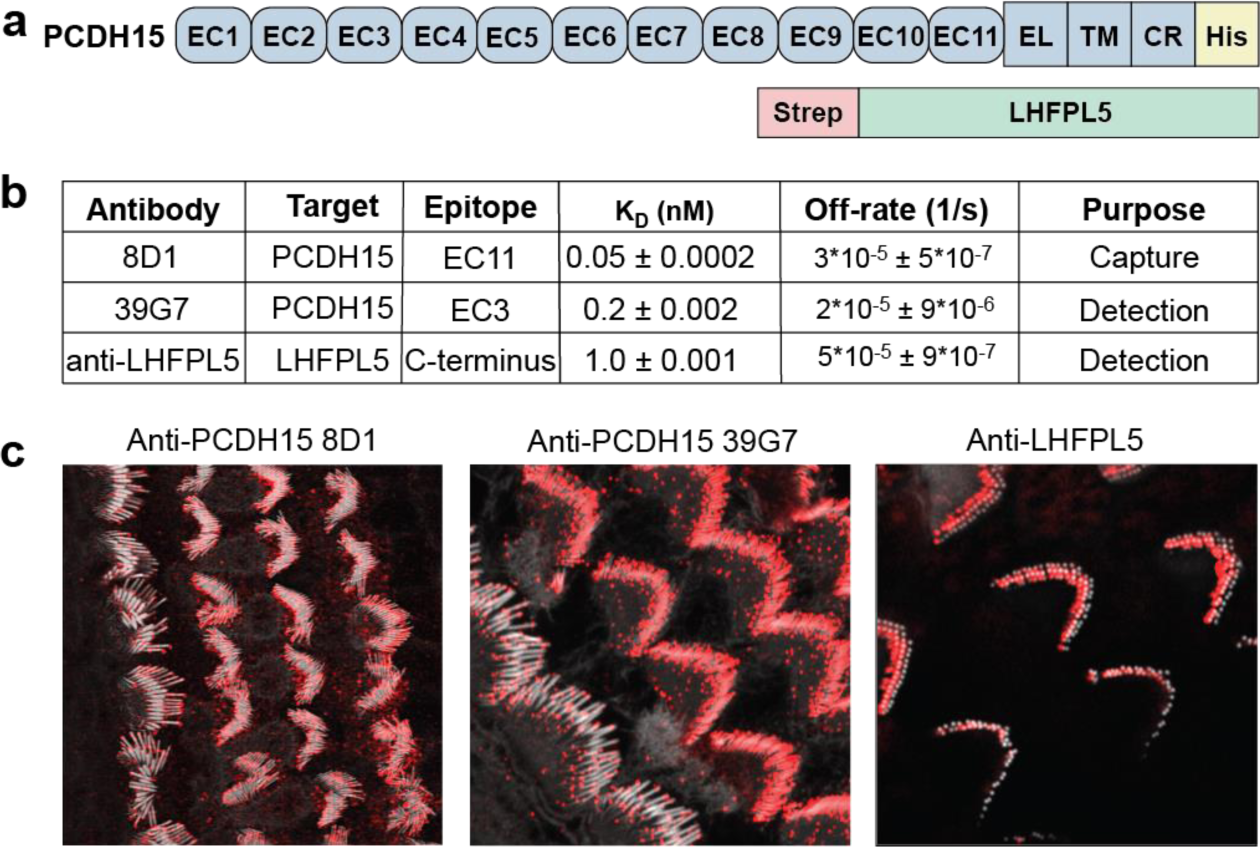
Characterization of reagents used for SiMPull and SiMoA assay development. **a,** Schematics of the recombinant PCDH15 and LHFPL5 constructs. **b,** Table of antibody binding constants and off-rates, measured by bio-layer interferometry. **c,** Immunostaining of WT cochlea with anti-PCDH15 and anti-LHFPL5 antibodies. Actin was stained with SirActin-405, shown in white, and mouse and rabbit monoclonal antibodies were detected with an anti-mouse-Alexa594 or anti-rabbit-Alexa594 secondary antibody, shown in red.

To develop a SiMPull assay of sufficient sensitivity to detect attomole quantities of PCDH15 and LHFPL5, we used recombinantly expressed PCDH15/LHFPL5 complex as a control. Multiple antibody combinations were assessed for capture and detection, allowing us to determine that 8D1 is the best antibody for capturing protein from low concentration samples, likely due to its fast on rate and slow dissociation rate (Supp. Fig. 1a). Passivated slides were coated with biotinylated 8D1 to capture PCDH15/LHFPL5, which was then detected with an anti-PCDH15 39G7 Fab fused to GFP or with an anti-LHFPL5 mAb conjugated to Alexa647 (Fig. 2a). We routinely were able to detect 3.2 attomoles of PCDH15 and 0.8 attomoles of LHFPL5 with signal-to-noise about 5-fold above background (Fig. 2b), indicating that the assay is adequately sensitive to detect native PCDH15 and LHFPL5. The difference in sensitivity between PCDH15 and LHFPL5 is likely due to the increased brightness of the Alexa647 fluorophore relative to a GFP molecule. For comparison, we performed a western blot with the same quantity of protein and discovered that we can detect only 12 picomoles of LHFPL5 with the anti-LHFPL5 mAb (Supp. Fig 2). 8D1 and 39G7 both recognize a folded, three-dimensional epitope and are unsuitable for western blot.

**Figure 2:**
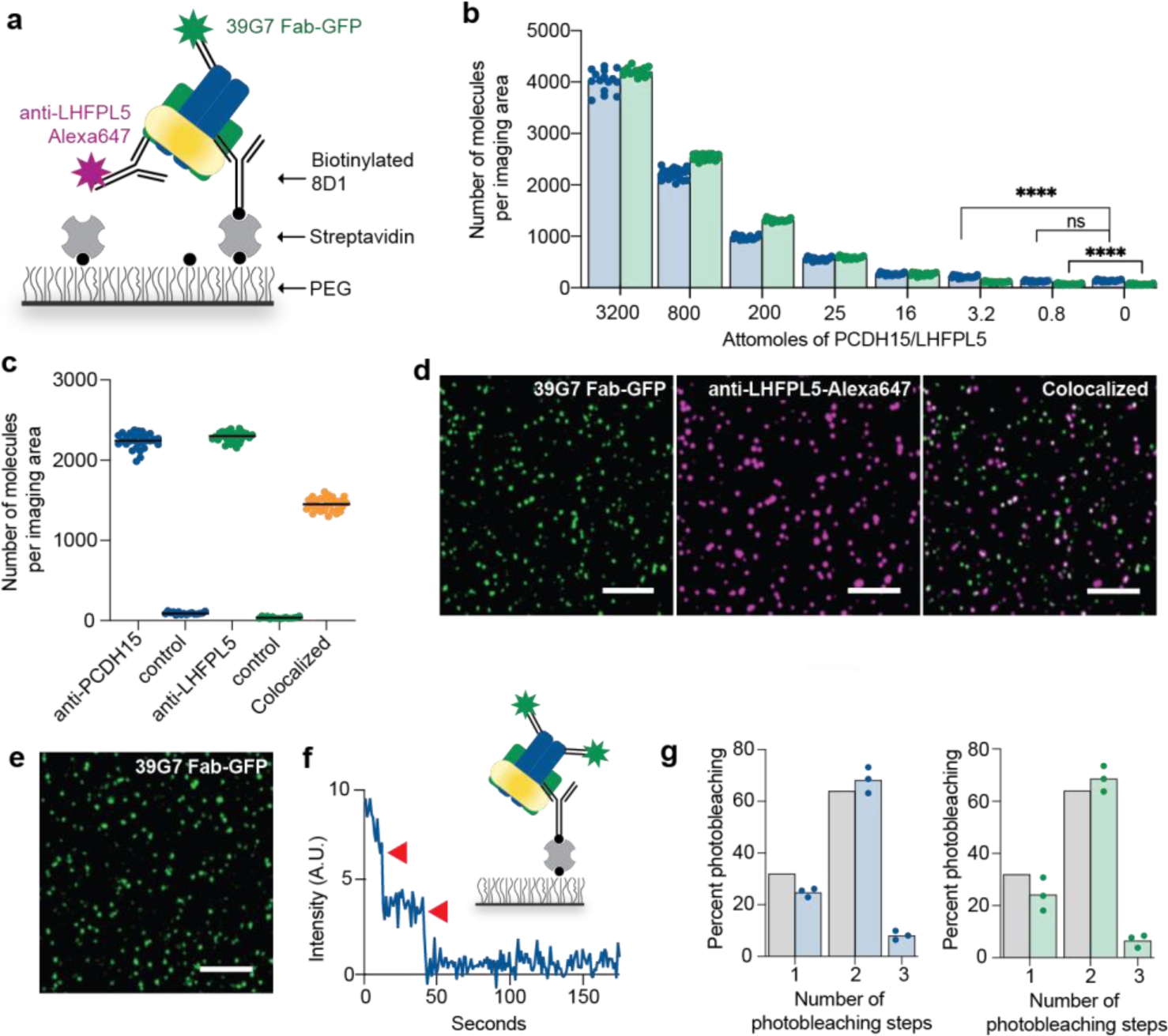
SiMPull assay development with recombinant PCDH15/LHFPL5. **a,** Schematic depiction of the SiMPull experiment. PCDH15/LHFPL5 complexes are immobilized on a passivated coverslip coated with anti-PCDH15 mAb 8D1. PCDH15 and LHFPL5 are detected with Fab-GFP or mAbs conjugated to Alexa dyes. **b,** SiMPull dilution analysis of PCDH15/LHFPL5 demonstrates that the assay can detect 10 attomoles of both PCDH15 and LHFPL5 when applied to the chamber in a volume of 10 µL. PCDH15 was detected with 39G7 Fab-GFP and LHFPL5 was detected with anti-LHFPL5 mAb conjugated to Alexa647 (**** p < 0.0001)**. c,** The total number of recombinant PCDH15, LHFPL5, and colocalized PCDH15/LHFPL5 molecules are shown. We observed 65% of PCDH15 molecules colocalized with LHFPL5 molecules. N= 30 images analyzed over two independent experiments. **d,** Representative TIRF images of recombinant PCDH15/LHFPL5 complex detected with anti-PCDH15 Fab 39G7-GFP and anti-LHFPL5 mAb. Scale bars are 5 µm. **e,** Representative TIRF image of PCDH15 detected with the 39G7 Fab-GFP. **f,** Representative trace showing two step photobleaching (red arrows) of the 39G7 Fab-GFP. Inset is a schematic of recombinant PCDH15/LHFPL5 complex detected with 39G7 Fab-GFP. **g,** Summary of photobleaching step distribution for PCDH15/LHFPL5 detected with 39G7 Fab-GFP (blue bars) or anti-LHFPL5 Fab-GFP (green bars). The photobleaching step distributions correspond to a binominal distribution that assumes a dimeric protein and 80% GFP maturation (grey bars). N= 600 spots were analyzed from three photobleaching movies (200 spots/movie) collected from two independent experiments.

Next, we analyzed the composition and stoichiometry of the recombinantly expressed PCDH15/LHFPL5 complex to confirm that it forms a heteromeric complex, as expected from structural studies ^9^. Application of 10 µL of 100 pM PCDH15/LHFPL5 complex to the sample chamber, followed by simultaneous application of the 39G7-GFP and anti-LHFPL5-Alexa647 antibodies, enabled assessment of the degree of colocalization between PCDH15 and LHFPL5. Recombinant LHFPL5 colocalizes with PCDH15 62% of the time (Fig. 2c), consistent with the expectation that the two proteins are found in the same complex. Although PCDH15 and LHFPL5 form a stable, homogenous complex, their lack of 100% colocalization can be attributed to a number of factors, including incomplete antibody binding, incomplete fluorophore labeling, and dissociation of the PCDH15/LHFPL5 complex.

To determine the stoichiometry of PCDH15 and LHFPL5, we captured the complex with 8D1 and, in separate experiments, labeled PCDH15 with 39G7 conjugated to GFP and labeled LHFPL5 with the anti-LHFPL5 Fab conjugated to GFP. The imaging area was photobleached for one minute and the resulting movies were analyzed to determine stoichiometry. Approximately 68% of 39G7 Fab-GFP molecules and 68% of anti-LHFPL5 Fab-GFP molecules bleached in two steps, demonstrating that PCDH15 and LHFPL5 are present in two copies each. These results demonstrate that we have developed a highly sensitive, robust assay that is capable of detecting and defining the stoichiometry of native PCDH15- and LHFPL5-containing complexes.

### Native PCDH15 and LHFPL5 form a heterotetrametric complex

While tremendous progress has been made in identifying the proteins that are necessary for MT in hair cells, the composition of the MT complex and organization of subunits within the complex is unknown. It is suspected that native PCDH15 forms a complex with LHFPL5 based on data from biochemical experiments and mutagenesis studies in mice ^7,9^. However, the interaction between native PCDH15 and LHFPL5 has never been demonstrated and it is unclear what portion of the PCDH15 population associates with LHFPL5.

To address these questions and demonstrate the utility of the SiMPull assay in detecting native MT subunits, we pulled down native PCDH15-containing complexes from mouse cochlea. Cochlea were homogenized in a buffer supplemented with a non-ionic detergent and the supernatant was applied to slides coated in the 8D1 antibody. Simultaneous application of the 39G7 Fab-GFP and anti-LHFPL5-Alexa647 mAb allowed us to measure colocalization between PCDH15 and LHFPL5. Approximately 58% of LHFPL5 subunits colocalized with PCDH15, indicating that the majority of PCDH15 molecules are in complex with LHFPL5 (Fig. 3a). Importantly, we did not observe a statistically significant signal above background for either PCDH15 or LHFPL5 when supernatant derived from the cochlea of PCDH15 knockout mice were applied to the slide. We similarly performed photobleaching experiments to assess the stoichiometry of native PCDH15 and LHFPL5. Approximately 59% of 39G7 Fab-GFP molecules and 63% of LHFPL5 molecules bleached in two-steps, indicating that both proteins are present in two copies. Our results indicate that PCDH15 and LHFPL5 form a heterotetrametric complex in hair cells and demonstrate that the SiMPull assay can be used to probe the assembly and stoichiometry of native MT subunits from solubilized mouse cochlea.

**Figure 3:**
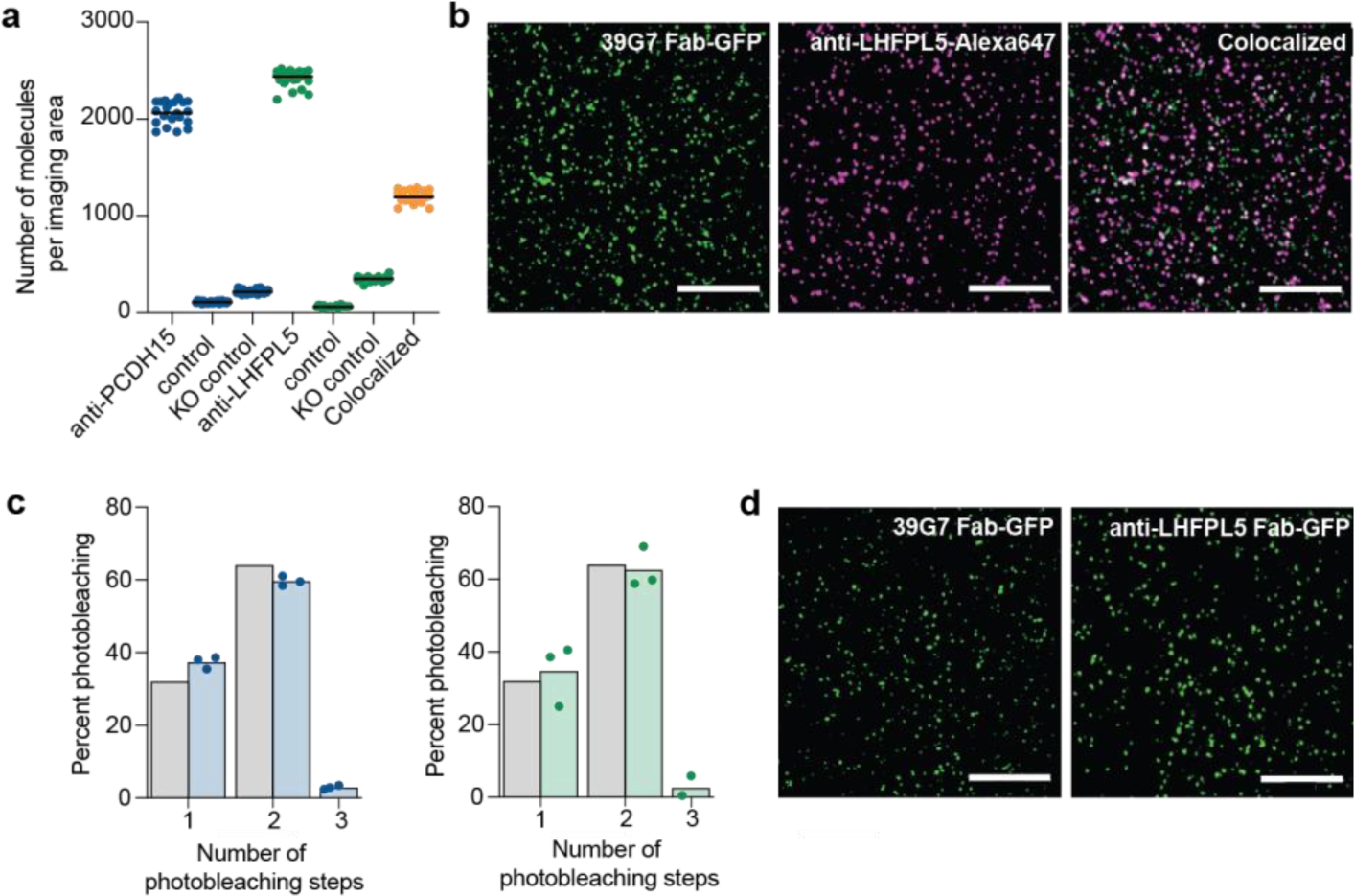
Native PCDH15/LHFPL5 form a heterotetrameric complex. **a,** Observed colocalization of native PCDH15 with native LHFPL5 was 58%. Negative controls included the standard control, the absence of 8D1 capture antibody, and PCDH15 KO cochlea. N= 25 images analyzed over two independent experiments. **b,** Summary of photobleaching step distribution of native PCDH15 (blue bars) and native LHFPL5 (green bars). The photobleaching step distributions correspond well with a binominal distribution (grey bars) that assumes a dimeric protein and 80% GFP maturation. N= 600 spots were analyzed from three photobleaching movies (200 spots/movie) collected from two independent experiments.

### Single molecule quantitation of native PCDH15 and LHFPL5

Many different approaches have been employed to estimate the number of MT subunits per stereocilia, all of which have relied on counting subunits in images of stereocilia. Freeze-etch electron microscopy images of stereocilia tips from bullfrogs and guinea pigs suggest that there is only one tip-link per stereocilia ^38^. Cryo-electron tomograms of stereocilia with gold nanoparticle-labeled PCDH15 indicate that there are 2, to more than 5, PCDH15 molecules per stereocilia tip and dozens more PCDH15 per shaft, amounting to greater than 15 attomoles per utricle. High resolution confocal microscopy images of fluorescently-labeled TMC1 similarly suggest that there are ∼8-20 TMC1 per stereocilia, or ∼24-60 attomoles of TMC1 per cochlea. While these methods have provided insight into the abundance of MT subunits, they are complicated by damage to the stereocilia during sample preparation and background staining of the labeled antibody. These methods are also labor intensive, requiring acquisition of dozens of high-resolution images. While SiMPull can be used to estimate the number of MT subunits, its accuracy is limited, in part due to the sample application method. The small volume of the SiMPull sample chamber requires iterative applications of the homogenized cochlea sample, introducing room for error. Further, the data from dilution experiments of recombinant PCDH15/LHFPL5 were not ideally linear (Fig. 2b). We therefore developed an ultrasensitive assay to accurately estimate the abundance of individual MT subunits in a mouse cochlea or utricle using single molecule array (SiMoA) ^36^.

In several respects, SiMoA is similar to SiMPull in that MT subunits in cell or tissue lysate are first captured on an antibody-coated surface, which for SiMoA is a paramagnetic bead, and then subsequently detected with a second antibody that is labeled with β-galactosidase (Fig. 4a). Following assembly of immunocomplexes on beads, the beads are loaded into an array of femtoliter-sized reaction wells, each of which can accommodate a single bead, and the β-galactosidase substrate is applied. Beads that possess an enzyme-labeled immunocomplex generate a relatively high local concentration of fluorescent product that is contained within the femtoliter reaction well, thus amplifying the signal and allowing for detection of single complexes.

**Figure 4:**
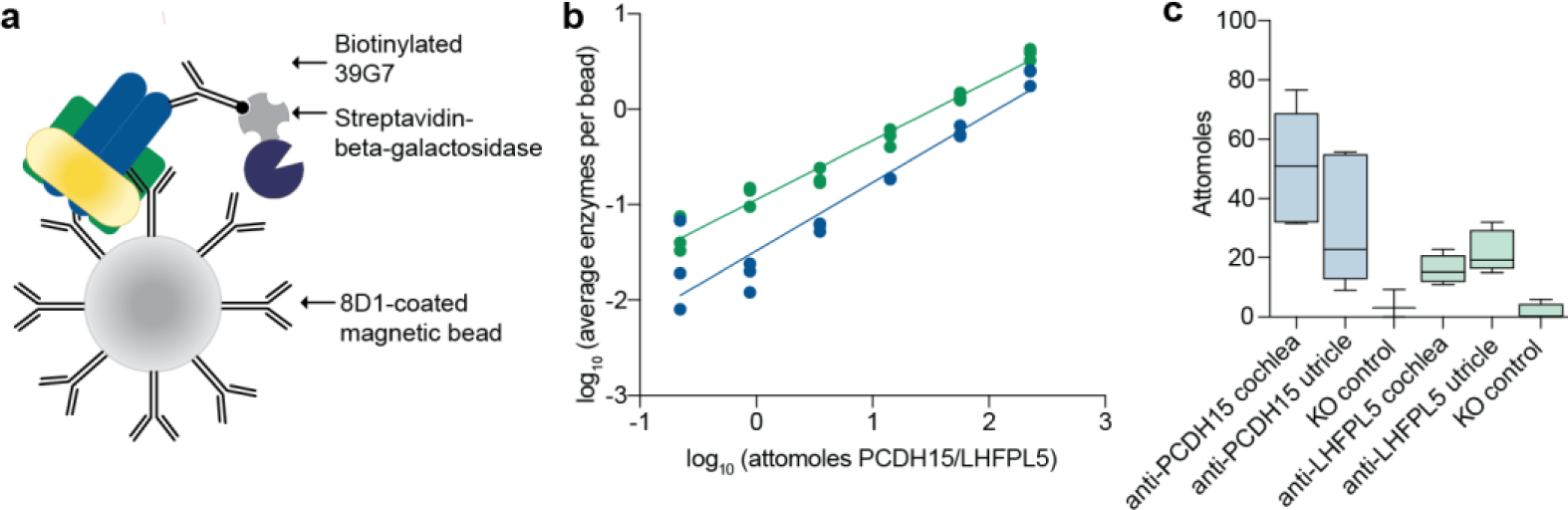
SiMoA assay development and quantification of native PCDH15 and the PCDH15/LHFPL5 complex. **a,** Schematic depiction of SiMoA capture and detection. PCDH15/LHFPL5 complexes are immobilized on a magnetic bead coated with anti-PCDH15 mAb 8D1. PCDH15 and LHFPL5 are detected with biotinylated anti-PCDH15 mAb 39G7 or anti-LHFPL5 mAb. Streptavidin conjugated to ß-galactosidase is used to quantify the number of PCDH15/LHFPL5 complexes per bead. **b,** Standard curve of recombinant PCDH15 (blue) and recombinant LHFPL5 (green). N=3 data points from three independent experiments are shown. **c,** A box plot depicting quantification of PCDH15 and LHFPL5 isolated from mouse cochlea or utricle. n = 6-8 cochlea or utricles for all samples.

To develop the SiMoA assay, we used recombinant PCDH15/LHFPL5 complex and the same antibodies that were used in SiMPull assays (Fig. 1b). Paramagnetic beads were functionalized with 8D1 and incubated either with recombinant protein sample or, following method optimization, cochlea supernatant that was prepared identically to SiMPull experiments. The beads were washed extensively and incubated with biotinylated 39G7 or anti-LHFPL5 mAbs, followed by a streptavidin-β-galactosidase conjugate (Fig. 4a). Using the SR-X Biomarker Detection System, the beads were isolated in arrays of 50 femtoliter reaction wells and the number of fluorescent wells were quantitated. We found that we were able to reliably detect 2 attomoles of PCDH15 and 4 attomoles of PCDH15/LHFPL5 using this assay, well within the range needed for detection of native MT subunits (Fig. 4b).

We applied the SiMoA assay to the quantification of native PCDH15 and LHFPL5 from cochlea and utricle supernatant (Fig. 4c). We found that there are an average of 55 attomoles of PCDH15 per cochlea and 25 attomoles per utricle. The individual measurements of PCDH15 abundance spanned a wide range, from 31-76 attomoles per cochlea and 9-55 attomoles per utricle, likely due to experimental variability, coupled with variations in mouse age. The number of PCDH15 molecules per stereocilia changes dramatically during hair bundle development up to post-natal day 9 (ref 39), so measurements of PCDH15 abundance will change due to small differences in mouse age. A signal above background was not detected for PCDH15 derived from cochlea of knockout animals. The number of PCDH15/LHFPL5 complexes, measured by detecting PCDH15-captured complexes with an anti-LHFPL5 antibody, fell within a similar range, 18 attomoles per cochlea and 21 attomoles per utricle. These data suggest that the majority of native PCDH15 is in complex with LHFPL5, in accordance with data obtained from the SiMPull experiments.

## Discussion

We have developed two single molecule assays for the detection, quantitation, and characterization of native MT subunits. We applied these assays to the native PCDH15/LHFPL5 complex isolated from mouse cochlea and utricles, demonstrating the exceptional sensitivity and utility of these assays for studying extremely low abundance proteins. The results of our SiMPull assay indicate that the majority of native PCDH15 is bound to LHFPL5 and the two proteins form a stable heterotetrameric complex. SiMoA quantitation reveals that there are an average of 18 attomoles of PCDH15/LHFPL5 complex and 55 attomoles of PCDH15 per cochlea, the first quantitative measurement of MT complex subunits.

The SiMPull and SiMoA assays have substantial potential for uncovering the molecular composition and organization of the MT complex, which have eluded scientists for decades. Immunoprecipitation and yeast two-hybrid experiments suggest multiple interactions between MT subunits, but these results are clouded by an inability to reconstitute the complex in heterologous cell lines. The SiMPull assay overcomes this barrier by isolating individual MT complexes from native cochlea and enabling analysis of their stoichiometry through photobleaching experiments, as well as their interactions with other subunits through colocalization experiments. These tools have the potential to elucidate the organization of the MT complex, as well as to report on the distribution of MT complexes if the subunit composition varies. Indeed, a recently-developed bead-based SiMPull assay indicates complex formation between TMC1 and LHFPL5, although colocalization and stoichiometry measurements were not possible due to the assay format ^40,41^. An additional application of SiMPull and SiMoA is the study of hair bundle assembly. Information regarding the developmental progression of protein expression, as well as data surrounding protein-protein interactions, are needed to understand how the tightly coordinated process of hair bundle assembly occurs ^42^.

There are, however, several limitations to these methods, the most importance of which is their reliance on high affinity antibodies. The assays cannot be employed if antibodies with good binding kinetics and affinities do not exist and there is no alternate way to capture the target protein, such as with a genetically-engineered tag. Additionally, it is possible that isolation of the complex results in the loss of weakly or transiently bound partners. Although we use gentle solubilization conditions, there is always a possibility that the removal of a protein from its native environment disrupts transient interactions. Further, both assays probe the composition of complexes derived from the entire cochlea, not just what is present at the stereocilia membrane. It is possible that the functional, fully-assembled MT complexes on stereocilia are only a fraction of the total complexes and that ‘differently assembled variants’ are present as intermediates or alternative species.

## Methods

### Construct design

The PCDH15 and LHFPL5 constructs correspond to the canonical *Mus musculus* sequences as recorded in the UniProt database. The PCDH15 construct was synthesized by Genscript and the LHFPL5 construct is the same construct used in previous studies ^9^. All constructs were cloned into the pEG BacMam vector under control of the CMV promoter, allowing expression by infection using Baculovirus produced in Sf9 cells. Details regarding the sequences, fluorophores, and affinity tags are described in Figure 1.

### Expression and purification of PCDH15 in complex with LHFPL5

The recombinant PCDH15/LHFPL5 complex was expressed and purified as described in Ge et al ^9^. Briefly, HEK293 tsa201 cells were co-infected with PCDH15 and LHFPL5 BacMam viruses at a MOI of 1:1. Cultures were supplemented with 10 mM sodium butyrate 12 hr post-infection and transferred to 30°C. Cells were harvested 48 hr post-infection and lysed in buffer containing 100 mM Tris pH 8.0, 150 mM NaCl, 1% (w/v) digitonin and protease inhibitors for 2 hr at 4°C. The solubilized material was incubated with Strep-Tactin resin, washed with buffer A containing 20 mM Tris pH 8.0, 150 mM NaCl, 0.07% (w/v) digitonin and eluted with buffer A plus 5 mM desthiobiotin to remove free His-tagged PCDH15. The elution was then incubated with TALON resin, washed with buffer A plus 10 mM imidazole to remove free strep-tagged LHFPL5, and further eluted with buffer A plus 200 mM imidazole. The complex was then further purified by size exclusion chromatography in buffer A. Peak fractions were collected and analyzed by SDS-PAGE.

### Expression and purification of antibody fragments

The anti-PCDH15 39G7 monoclonal antibody was generated as described in Elferich et al. ^31^. The anti-LHFPL5 monoclonal antibody was obtained from Abcam and the amino acid sequence of the variable domain was determined by mass spectrometry. The anti-LHFPL5 construct was synthesized by Genscript. The DNA sequences encoding the heavy and light chains of the variable domains of the 39G7 and anti-LHFPL5 antibodies were cloned into the pEG BacMam vector for baculovirus expression in HEK293 tsa201 cells. An mVenus tag and 8xHis tag were added to the C-terminus of the heavy chain in both constructs. Cells were infected with 39G7 or anti-LHFPL5 BacMam viruses at a MOI of 1:1. Cultures were supplemented with 10 mM sodium butyrate 12 hr post-infection and transferred to 30°C. Cells were harvested 96 hr post-infection and the cell media was filtered and concentrated to 200 mL. Concentrated media was incubated with TALON resin, washed with TBS plus 10 mM imidazole, and eluted with TBS plus 200 mM imidazole. The antibody fragments were further purified using size exclusion chromatography in PBS. The affinities of the 8D1, 39G7 and anti-LHFPL5 monoclonal antibodies for their antigen were determined with biolayer interferometry using an OctetRED384 instrument.

### Cochlea and utricle solubilization

Cochlea and utricle samples were prepared by homogenizing cochlea or utricles from post-natal day 6 mice on ice with a pestle homogenizer in lysis buffer consisting of 1% C12M, 0.2% CHS, 50 mM Tris, 40 mM NaCl, 10 mM KCl, 1 mM EDTA, 1 mM protease inhibitors (0.8 µM aprotinin, 2 µg/ml leupeptin and 2 µM pepstatin), 0.2 mg/mL BSA, pH 8.0. 20 μL of lysis buffer was added per cochlea. Homogenized cochlea were incubated for 1 hr at 4 °C and insoluble material was pelleted by centrifuging for 10 minutes at 14,000 rpm. The supernatant was collected and used immediately for SiMPull or SiMoA experiments.

### Single molecule pulldown

Coverslips and glass slides were prepared as described in Jain et al ^37^. In brief, the coverslips and glass slides were extensively cleaned, passivated, and coated with methoxy polyethylene glycol (mPEG) and 2% biotinylated PEG. A flow chamber was created by drilling 0.75 mm holes in the quartz slide and placing double-sided tape between the holes. A coverslip was placed on top of the slide and the edges were sealed with epoxy, creating tiny flow chambers. 0.25 mg/mL streptavidin was then applied to the slide, allowed to incubate for 5 minutes, and washed off with T50 BSA buffer consisting of 50 mM Tris pH 8.0, 50 mM NaCl, 0.25 mg/mL BSA. Anti-PCDH15 8D1 capture antibody was biotinylated using NHS-PEG4-biotin (Thermo Fisher CAS# 21330) and anti-LHFPL5 antibody was labeled with Alexa647 using a Zip Alexa Fluor Rapid Antibody Labeling Kit (Thermo Fisher CAS#Z11235). Excess dye and biotin were removed using two Zeba spin desalting columns per labeling reaction and labeling was confirmed using fluorescence-detection size exclusion chromatography (FSEC) ^43^. Biotinylated 8D1 at approximately 7 μg/mL was applied to the slide, allowed to incubate for 10 minutes, and washed off with buffer B containing 0.005% C12M, 0.001% CHS, 50 mM Tris, 40 mM NaCl, 10 mM KCl, 0.1 mM EDTA, 0.2 mg/mL BSA, pH 8.0. For recombinant PCDH15/LHFPL5 complex, the sample was diluted 50 pM in buffer B and incubated in the chamber for 5 minutes. For cochlea supernatant, the sample was applied to the chamber in 10 μL increments, with a 5 minute incubation per application. Following sample application, the chamber was washed and 3 μg/mL fluorophore-labeled detection antibody was applied to the chamber for 5 minutes. The chamber was imaged immediately after the final wash step using total internal reflection (TIRF) microscopy and 15 images at different locations were collected per chamber. Photobleaching movies were collected by exposing the imaging area for 2 minutes.

Molecule quantitation and colocalization were determined using the ComDet v0.5.5 plugin for FIJI ^44^, available at (https://github.com/UU-cellbiology/ComDet). Photobleaching movies were analyzed using a custom python script that is available on Zenodo (https://zenodo.org/records/8161179).

### Single molecule array

Recombinant PCDH15/LHFPL5 complex and cochlea samples were quantitated using the Quanterix homebrew assay kit and a Quanterix SR-X Biomarker Detection System. Anti-PCDH15 8D1 was conjugated to paramagnetic beads for capturing analyte and anti-PCDH15 39G7 mAb or anti-LHFPL5 mAb was biotinylated for analyte detection according to instructions in the homebrew assay kit. The 2-step method was employed wherein beads, sample, and detection antibody were incubated simultaneously for 30 minutes with continuous shaking at room temperature. Sample binding buffer consisted of 1% C12M, 0.2% CHS, 1% casein in PBS, 1 mM EDTA, protease inhibitors, and 10 μg/mL mouse IgG. After incubation, the beads were washed with buffer C containing 0.005% C12M in PBS. 120 pM streptavidin beta-galactosidase (SBG) was applied for 7 minutes and beads were washed again with buffer C, followed by a short wash with SiMoA wash buffer B (buffer composition is proprietary), and allowed to dry for 10 minutes. Dried plates were then placed in the SR-X for substrate application and analysis. All samples were run in duplicate.

## Acknowledgements

The authors would like to thank members of the Gouaux and Baconguis laboratories for helpful discussions and Rachel Courtney for proof reading the manuscript. The authors would also like to acknowledge the staff in the OHSU Advanced Light Microscopy Core (RRID:SCR_009961), especially Stefanie Petrie, for help with cochlea imaging. This work was supported by NIH grant 1F32DC017894 to S.C. E.G. gratefully acknowledges J. LaCroute and B. LaCroute for support, and is an investigator of the HHMI. T.H. is an investigator of the HHMI.

## Author Contributions

S.C., J.M., and J.E. performed the experiments. All authors contributed to the design, analysis, and interpretation of experimental data. S.C. and E.G. wrote the manuscript and all authors contributed to the review and editing of the manuscript.

## Data Availability

Custom code used for analyzing single-molecule photobleaching trajectories in this study is available at Zenodo (https://doi.org/10.5281/zenodo.8161179).

---

**Supplementary Figure 1:**
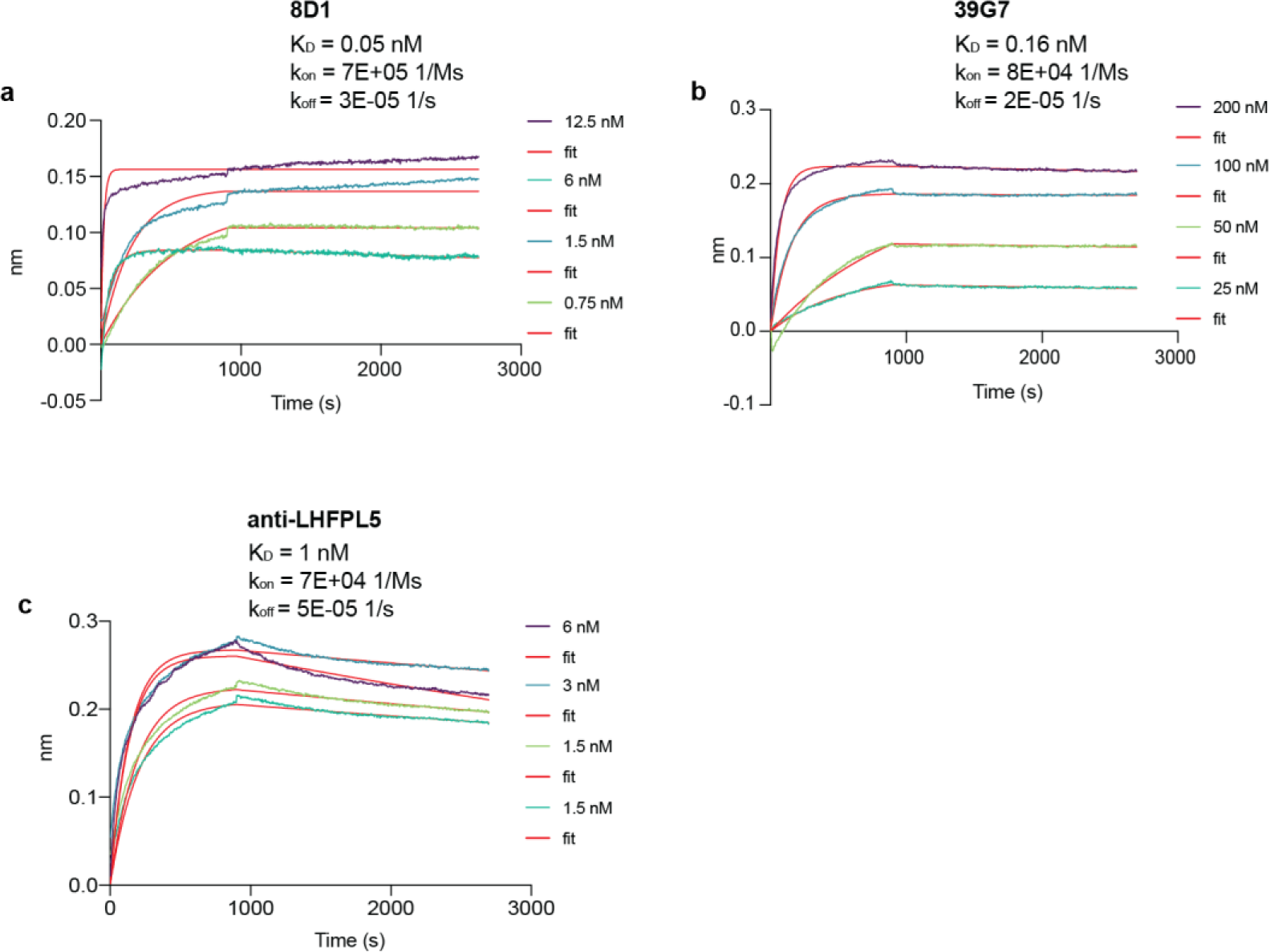
Bio-layer interferometry measurements of anti-PCDH15 and anti-LHFPL5 antibodies. Experimental traces are shown for anti-PCDH15 8D1 **(a),** anti-PCDH15 39G7 **(b),** and anti-LHFPL5 **(c).** Concentrations of monoclonal antibody ranged from 0.75 – 200 nM depending on the antibody.

**Supplementary Figure 2:**
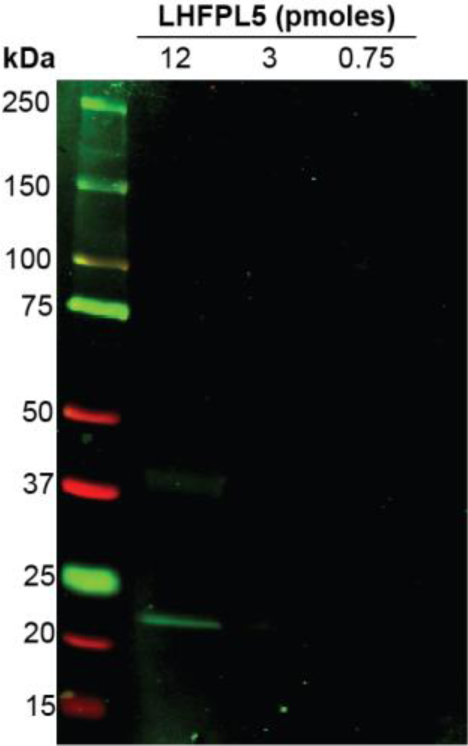
Western blot of recombinant LHFPL5. Recombinant LHFPL5 was probed with anti-LHFPL5 monoclonal antibody at 0.75 pM, 3 pM, and 12 pM, to facilitate direct comparison to SiMPull and SiMoA experiments.

## References

1 Zheng, W. & Holt, J. R. The Mechanosensory Transduction Machinery in Inner Ear Hair Cells. Annu Rev Biophys 50, 31–51 (2021). 10.1146/annurev-biophys-062420-081842

2 Kazmierczak, P. et al. Cadherin 23 and protocadherin 15 interact to form tip-link filaments in sensory hair cells. Nature 449, 87–91 (2007). 10.1038/nature06091

3 Sakaguchi, H., Tokita, J., Muller, U. & Kachar, B. Tip links in hair cells: molecular composition and role in hearing loss. Curr Opin Otolaryngol Head Neck Surg 17, 388–393 (2009). 10.1097/MOO.0b013e3283303472

4 Pan, B. et al. TMC1 and TMC2 are components of the mechanotransduction channel in hair cells of the mammalian inner ear. Neuron 79, 504–515 (2013). 10.1016/j.neuron.2013.06.019

5 Pan, B. et al. TMC1 Forms the Pore of Mechanosensory Transduction Channels in Vertebrate Inner Ear Hair Cells. Neuron 99, 736–753 e736 (2018). 10.1016/j.neuron.2018.07.033

6 Jia, Y. et al. TMC1 and TMC2 Proteins Are Pore-Forming Subunits of Mechanosensitive Ion Channels. Neuron 105, 310–321 e313 (2020). 10.1016/j.neuron.2019.10.017

7 Xiong, W. et al. TMHS is an integral component of the mechanotransduction machinery of cochlear hair cells. Cell 151, 1283–1295 (2012). 10.1016/j.cell.2012.10.041

8 Beurg, M., Xiong, W., Zhao, B., Muller, U. & Fettiplace, R. Subunit determination of the conductance of hair-cell mechanotransducer channels. Proc Natl Acad Sci U S A 112, 1589–1594 (2015). 10.1073/pnas.1420906112

9 Ge, J. et al. Structure of mouse protocadherin 15 of the stereocilia tip link in complex with LHFPL5. Elife 7 (2018). 10.7554/eLife.38770

10 Zhao, B. et al. TMIE is an essential component of the mechanotransduction machinery of cochlear hair cells. Neuron 84, 954–967 (2014). 10.1016/j.neuron.2014.10.041

11 Pacentine, I. V. & Nicolson, T. Subunits of the mechano-electrical transduction channel, Tmc1/2b, require Tmie to localize in zebrafish sensory hair cells. PLoS Genet 15, e1007635 (2019). 10.1371/journal.pgen.1007635

12 Cunningham, C. L. et al. TMIE Defines Pore and Gating Properties of the Mechanotransduction Channel of Mammalian Cochlear Hair Cells. Neuron 107, 126–143 e128 (2020). 10.1016/j.neuron.2020.03.033

13 Giese, A. P. J. et al. CIB2 interacts with TMC1 and TMC2 and is essential for mechanotransduction in auditory hair cells. Nat Commun 8, 43 (2017). 10.1038/s41467-017-00061-1

14 Liang, X. et al. CIB2 and CIB3 are auxiliary subunits of the mechanotransduction channel of hair cells. Neuron 109, 2131–2149 e2115 (2021). 10.1016/j.neuron.2021.05.007

15 Cunningham, C. L. et al. The murine catecholamine methyltransferase mTOMT is essential for mechanotransduction by cochlear hair cells. Elife 6 (2017). 10.7554/eLife.24318

16 Erickson, T. et al. Integration of Tmc1/2 into the mechanotransduction complex in zebrafish hair cells is regulated by Transmembrane O-methyltransferase (Tomt). Elife 6 (2017). 10.7554/eLife.28474

17 Michel, V. et al. Interaction of protocadherin-15 with the scaffold protein whirlin supports its anchoring of hair-bundle lateral links in cochlear hair cells. Sci Rep 10, 16430 (2020). 10.1038/s41598-020-73158-1

18 Delprat, B. et al. Myosin XVa and whirlin, two deafness gene products required for hair bundle growth, are located at the stereocilia tips and interact directly. Hum Mol Genet 14, 401–410 (2005). 10.1093/hmg/ddi036

19 Belyantseva, I. A. et al. Myosin-XVa is required for tip localization of whirlin and differential elongation of hair-cell stereocilia. Nat Cell Biol 7, 148–156 (2005). 10.1038/ncb1219

20 Belyantseva, I. A., Boger, E. T. & Friedman, T. B. Myosin XVa localizes to the tips of inner ear sensory cell stereocilia and is essential for staircase formation of the hair bundle. Proc Natl Acad Sci U S A 100, 13958–13963 (2003). 10.1073/pnas.2334417100

21 Sotomayor, M., Weihofen, W. A., Gaudet, R. & Corey, D. P. Structure of a force-conveying cadherin bond essential for inner-ear mechanotransduction. Nature 492, 128–132 (2012). 10.1038/nature11590

22 Giese, A. P. J. et al. Complexes of vertebrate TMC1/2 and CIB2/3 proteins form hair-cell mechanotransduction cation channels. bioRxiv (2023). 10.1101/2023.05.26.542533

23 Jeong, H. et al. Structures of the TMC-1 complex illuminate mechanosensory transduction. Nature 610, 796–803 (2022). 10.1038/s41586-022-05314-8

24 Maeda, R. et al. Tip-link protein protocadherin 15 interacts with transmembrane channel-like proteins TMC1 and TMC2. Proc Natl Acad Sci U S A 111, 12907–12912 (2014). 10.1073/pnas.1402152111

25 Qiu, X., Liang, X., Llongueras, J. P., Cunningham, C. & Muller, U. The tetraspan LHFPL5 is critical to establish maximal force sensitivity of the mechanotransduction channel of cochlear hair cells. Cell Rep 42, 112245 (2023). 10.1016/j.celrep.2023.112245

26 Soler, D. C. et al. An uncharacterized region within the N-terminus of mouse TMC1 precludes trafficking to plasma membrane in a heterologous cell line. Sci Rep 9, 15263 (2019). 10.1038/s41598-019-51336-0

27 Beurg, M., Evans, M. G., Hackney, C. M. & Fettiplace, R. A large-conductance calcium-selective mechanotransducer channel in mammalian cochlear hair cells. J Neurosci 26, 10992–11000 (2006). 10.1523/JNEUROSCI.2188-06.2006

28 Beurg, M., Fettiplace, R., Nam, J. H. & Ricci, A. J. Localization of inner hair cell mechanotransducer channels using high-speed calcium imaging. Nat Neurosci 12, 553–558 (2009). 10.1038/nn.2295

29 Denk, W., Holt, J. R., Shepherd, G. M. & Corey, D. P. Calcium imaging of single stereocilia in hair cells: localization of transduction channels at both ends of tip links. Neuron 15, 1311–1321 (1995). 10.1016/0896-6273(95)90010-1

30 Beurg, M. et al. Variable number of TMC1-dependent mechanotransducer channels underlie tonotopic conductance gradients in the cochlea. Nat Commun 9, 2185 (2018). 10.1038/s41467-018-04589-8

31 Elferich, J. et al. Molecular structures and conformations of protocadherin-15 and its complexes on stereocilia elucidated by cryo-electron tomography. Elife 10 (2021). 10.7554/eLife.74512

32 Iha, K. et al. Ultrasensitive ELISA Developed for Diagnosis. Diagnostics (Basel) 9 (2019). 10.3390/diagnostics9030078

33 He, X. & Patfield, S. A. Immuno-PCR Assay for Sensitive Detection of Proteins in Real Time. Methods Mol Biol 1318, 139–148 (2015). 10.1007/978-1-4939-2742-5_14

34 Niemeyer, C. M., Adler, M. & Wacker, R. Immuno-PCR: high sensitivity detection of proteins by nucleic acid amplification. Trends Biotechnol 23, 208–216 (2005). 10.1016/j.tibtech.2005.02.006

35 Jain, A. et al. Probing cellular protein complexes using single-molecule pull-down. Nature 473, 484–488 (2011). 10.1038/nature10016

36 Rissin, D. M. et al. Single-molecule enzyme-linked immunosorbent assay detects serum proteins at subfemtomolar concentrations. Nat Biotechnol 28, 595–599 (2010). 10.1038/nbt.1641

37 Jain, A., Liu, R., Xiang, Y. K. & Ha, T. Single-molecule pull-down for studying protein interactions. Nat Protoc 7, 445–452 (2012). 10.1038/nprot.2011.452

38 Kachar, B., Parakkal, M., Kurc, M., Zhao, Y. & Gillespie, P. G. High-resolution structure of hair-cell tip links. Proc Natl Acad Sci U S A 97, 13336–13341 (2000). 10.1073/pnas.97.24.13336

39 Goodyear, R. J., Marcotti, W., Kros, C. J. & Richardson, G. P. Development and properties of stereociliary link types in hair cells of the mouse cochlea. J Comp Neurol 485, 75–85 (2005). 10.1002/cne.20513

40 Yu, X. et al. Deafness mutation D572N of TMC1 destabilizes TMC1 expression by disrupting LHFPL5 binding. Proc Natl Acad Sci U S A 117, 29894–29903 (2020). 10.1073/pnas.2011147117

41 Zhao, Q. et al. Analyzing protein-protein interactions in rare cells using microbead-based single-molecule pulldown assay. Lab Chip 21, 3137–3149 (2021). 10.1039/d1lc00260k

42 Barr-Gillespie, P. G. Assembly of hair bundles, an amazing problem for cell biology. Mol Biol Cell 26, 2727–2732 (2015). 10.1091/mbc.E14-04-0940

43 Goehring, A. et al. Screening and large-scale expression of membrane proteins in mammalian cells for structural studies. Nat Protoc 9, 2574–2585 (2014). 10.1038/nprot.2014.173

44 Schindelin, J., et al. Fiji: an open-source platform for biological-image analysis. Nat Methods 9, 676–682 (2012). 10.1038/nmeth.2019

